# The mRNACalc web server accounts for the hypochromicity of modified nucleosides and enables the accurate quantification of nucleoside-modified mRNA

**DOI:** 10.1101/2023.07.27.550903

**Authors:** Esteban Finol, Sarah E. Krul, Sean J. Hoehn, Carlos E. Crespo-Hernández

## Abstract

Nucleoside-modified mRNA technologies necessarily incorporate N^1^-methylpseudouridine into the mRNA molecules to prevent over-stimulation of cytoplasmic RNA sensors. Despite this modification, mRNA concentrations remain mostly determined through measurement of UV absorbance at 260 nm wavelength (A_260_). Herein, we report that the N^1^-methylpseudouridine absorbs approximately 40% less UV light at 260 nm than uridine, and its incorporation into mRNAs leads to the under-estimation of nucleoside-modified mRNA concentrations, with 5-15% error, in a mRNA sequence dependent manner. We therefore examined the RNA quantification methods and developed the mRNACalc web server. It accounts for the molar absorption coefficient of modified nucleotides at 260 nm wavelength, the RNA composition of the mRNA, and the A_260_ of the mRNA sample to enable accurate quantification of nucleoside-modified mRNAs. The webserver is freely available at https://www.mrnacalc.com.

**Graphical Abstract:** 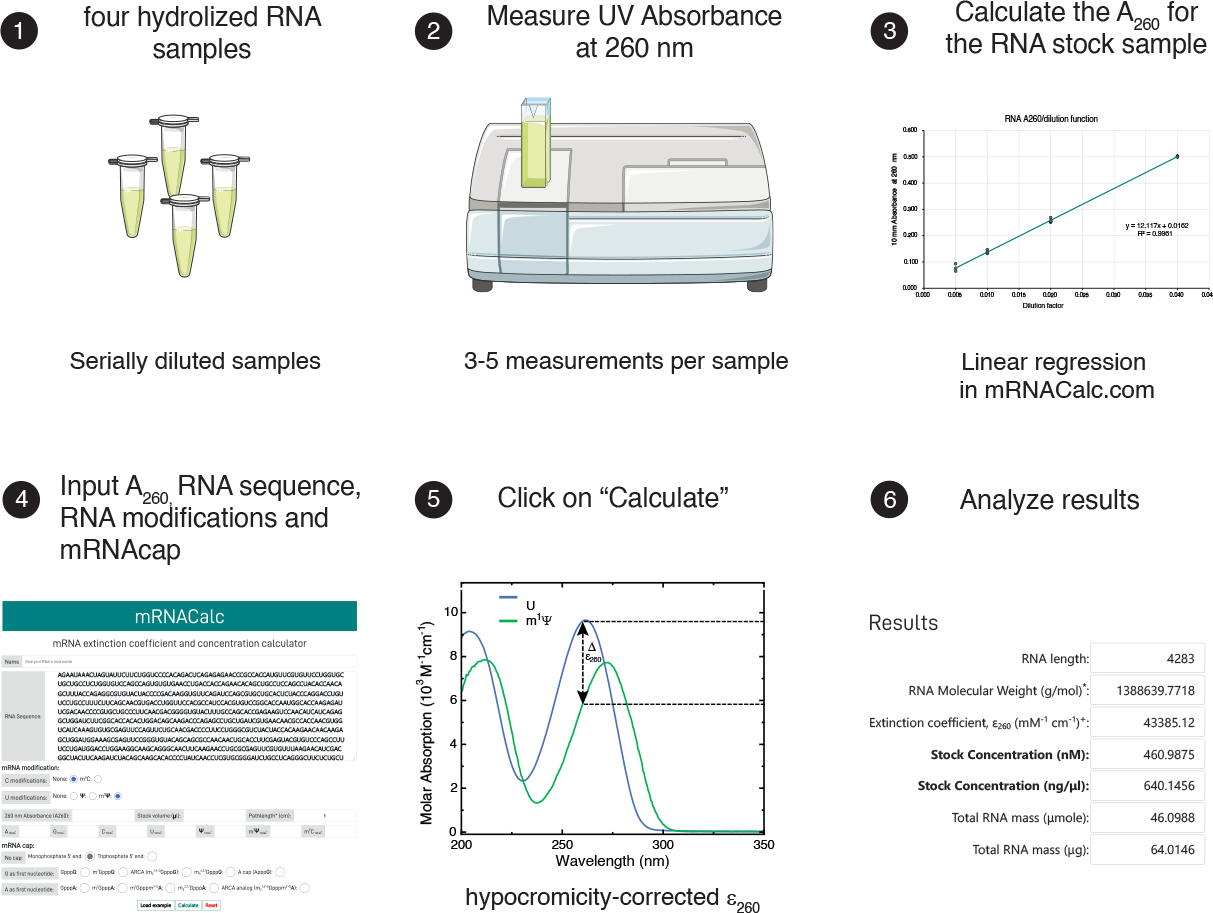

## Introduction

The therapeutic use of messenger RNA (mRNA) has sparked great optimism in the development of novel vaccines and therapeutics against a myriad of infectious or yet incurable diseases (1). The mRNA technology enables the production of antigenic, functional, and/or therapeutic proteins by introducing mRNA into the human body and cells (2). Since mRNAs act in the cytoplasm transiently, they do not bear any risk of integration into the host cell genome. Most importantly, the mRNA technology enables rapid, cost-efficient and scalable production, which is free of cellular (cell cultures) or animal materials (3). Thus, mRNA technologies facilitate manufacturing and allow for a rapid response to emerging infectious diseases, as emphatically underscored by the rapid rollout of COVID-19 mRNA vaccines in many parts of the world. Modified nucleosides, such as pseudouridine (Ψ), N^1^-methylpseudouridine (m^1^Ψ) and 5-methylcytidine (m^5^C), are often incorporated into the mRNA molecules. Such modifications reduce stimulation of cytoplasmic RNA sensors, such as toll-like receptor 3 and 7, for improved safety profiles and enhanced mRNA translation (4, 5). However, how modified nucleosides affect mRNA concentration measurements and potentially confound pre-clinical dosing, efficacy, and toxicology studies, which could make or break further clinical development of any therapeutic, remains undefined.

The determination of RNA concentration often relies on measurements of its UV absorbance at 260 nm wavelength (A_260_) and the implementation of the Beer–Lambert law (6). The accuracy of these measurements is scattered by the variable hypochromicity of RNA due to its sequence-dependent folding. The molar absorption coefficient (MAC or extinction coefficient, ε) of a folded RNA at 260 nm (ε_260_) is reduced as compared to its unfolded state (7). This difference is buffer- and concentration-dependent and arises from changes in the chemical environment of the nucleobases – the main chromophore, due to base-pairing, stacking, intermolecular interactions, and other conformational changes. Considering these variabilities, a rough estimation for the ε_260_ of any single stranded RNA (40 μg/ml per absorbance unit) is extensively used and its associated ±10–20 % error in the estimation of RNA concentration is widely accepted (6). This error range may suffice to assess dose-response for mRNA therapeutics across several orders of magnitude *in celula* or *in vivo* experiments. Yet it would be valuable to know concentrations at higher accuracy for the development of mRNA technologies. Our particular concern is in measurements of self-amplifying RNAs (saRNA) and nucleoside-modified mRNAs. The logarithmic amplification of saRNA can convert a 20 % accepted error in RNA concentration into a several fold differences in dose-response between one experiment and the subsequent replicates. The chemical modifications on the nucleobases of mRNA can also induce profound changes in the mRNA MAC, hindering the accurate quantification of nucleoside-modified mRNA concentrations.

To attain greater accuracy in RNA quantification, RNA molecules are hydrolysed prior to UV absorbance determination using a combination of thermal and alkaline hydrolysis (6, 8). The RNA hydrolysis shifts the hypochromic folded state of the RNA to the hyperchromic state of the single monophosphate nucleotides (9). Since the precise MAC of the four standard nucleotides in aqueous buffered solution are known, the molar absorption of any hydrolysed mRNA can be calculated as the sum of the molar absorption of its nucleotide compositions. Thus, upon the determination of the 260 nm UV absorbance (A_260_), the RNA concentration can be quantified with an error of ∼ 4 % using these methods (6). The incorporation of modified nucleosides can alter the RNA molar absorption and increase the error of the measurements in an RNA sequence-dependent manner. Other non-UV-spectroscopic methods relying on the unspecific RNA binding of fluorophores (such as RiboGreen, Thermo Fisher Scientific) for the determination of RNA concentration may help to overcome any change in the MAC of modified nucleoside mRNA. However, the impact of RNA modifications on the binding affinity of these fluorophores also remains unknown.

Herein, we report our effort to revisit and determine the molar absorption coefficients of modified nucleosides (Ψ, m^1^Ψ and m^5^C). We also examined three different methods for RNA hydrolysis and provided them along with the mRNACalc web server. This web tool incorporates the most recently revised ε_260_ for standard, modified and mRNA capping nucleosides, allowing the accurate determination of standard and nucleoside-modified mRNAs using UV spectroscopy. Once the RNA sequence, the A_260_ and the RNA stock volume values are provided as input, the mRNACalc web server calculates the RNA stock concentration in nM and ng/μl and the total RNA mass in μmole and μg.

## Results and discussion

To assess the impact of chemical modifications on the spectrophotometric parameters of nucleosides for mRNA quantification, we examined the modified nucleosides that have recently been employed in nucleoside-modified mRNA technologies: pseudouridine (Ψ), N^1^-methyl-pseudouridine (m^1^Ψ), and 5-methylcytidine (m^5^C). Pseudouridine is an isomer of uridine – the standard nucleoside in RNA. Pseudouridine, as opposed to other nucleosides, is a carbon-carbon ribofuranosyl nucleoside, i.e., the uracil nucleobase is linked to the ribose through its fifth carbon, instead of a N^1^-linkage (10). This unique arrangement places the N^1^-imino group toward the so-called “C-H” edge of the pyrimidine ring and confers additional properties to this edge in pseudouridine. This imino hydrogen proton is susceptible to hydrogen bonding, chemical exchange, and chemical modifications such as N^1^-methylation. Thus, the N^1^-methyl-pseudouridine, as well as the m^5^C, represents a modification of the C-H edge of the pyrimidine nucleobase. A similar 5-methyl modification also differentiates uridine from thymidine. The influence of a 5-methyl substituent on the UV molar absorption of pyrimidine rings is well known since the 1940’s when Sister Miriam Michael Stimson showed that it provokes a subtle reduction in molar absorbance and a shift of the peak maximum (ε_max_ and λ_max,_ respectively) to a longer wavelength (lower energy) – the so-called bathochromic or red shift (11–14). This red shift also leads to a substantial ε_260_ reduction for the methylated pyrimidine nucleosides. For the uridine to thymidine and cytidine to 5-methylcytidine comparisons, the respective peak shifts (Δλ_max_) are + 5 and + 7 nm with a 11.4 % and 20.8 % reduction in their ε_260_ (Fig.1a and 1b). For the Ψ and m^1^Ψ curves, we have observed a similar bathochromic shift (Δλ_max_ = + 9 nm, Fig. 1c), leading to a reduced molar absorption at 260 nm for m^1^Ψ (Δε_260_ = - 22.8 %). More importantly, m^1^Ψ is hypochromic as compared to uridine at λ_max_ (Δε_max_ = - 21 %) and, due to the bathochromic shift, m^1^Ψ absorbs 39.8 % less than uridine at 260 nm (Fig. 1d). Thus, the substitution of uridine by m^1^Ψ in mRNA technologies can lead to substantial changes in the spectrophotometric properties of the mRNA and may lead to the underestimation of nucleoside-modified mRNA concentrations.

**Fig. 1:**
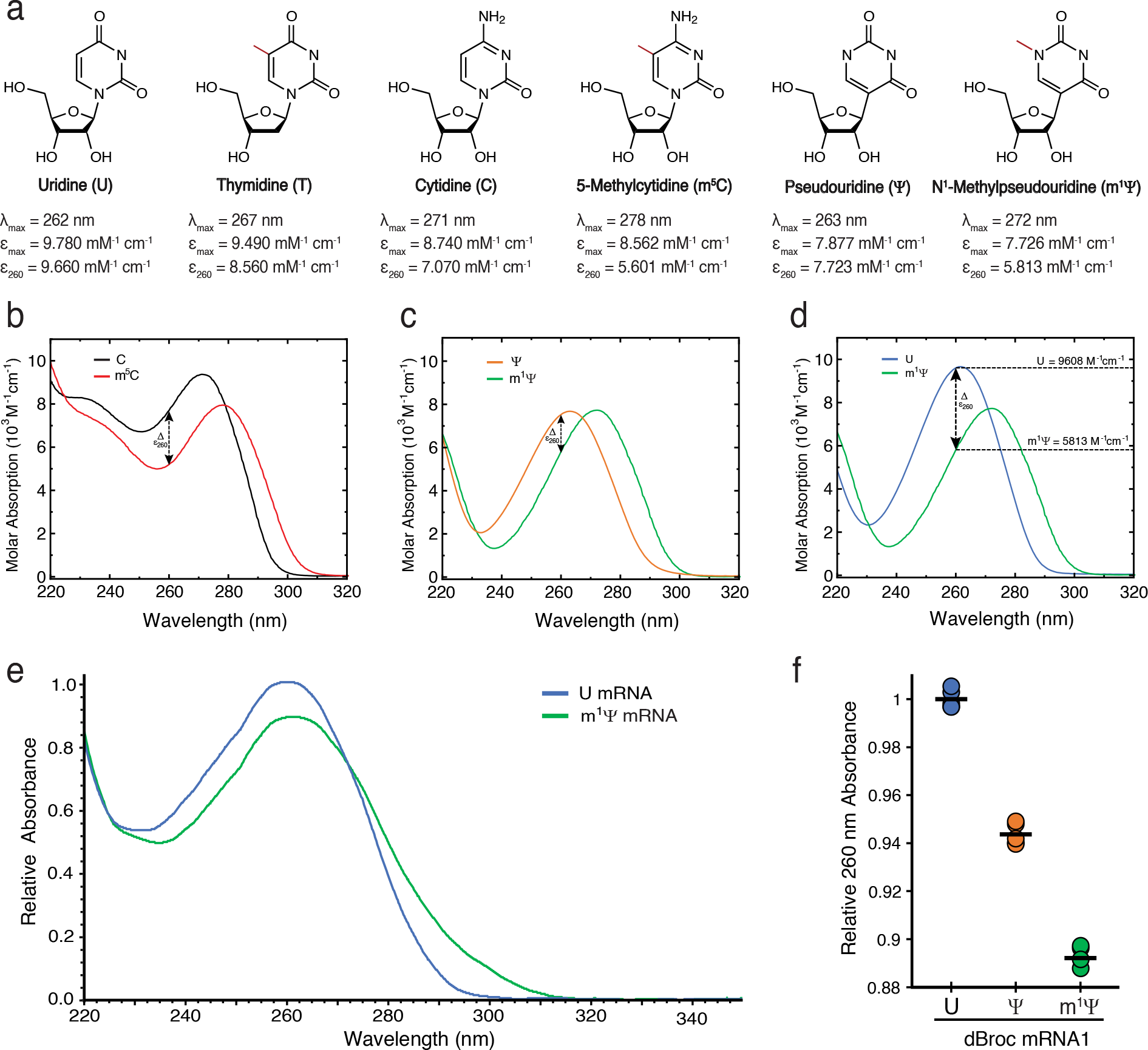
The nucleobase methylation and its bathochromic effect on the UV molar absorption spectra of pyrimidines and nucleoside-modified mRNAs. **a**, Skeletal formula of uridine, thymidine, cytidine, 5-methylcytidine, pseudouridine and N1-methylpseudouridine. The methyl substituents are highlighted in red. These λ_max_, ε_max_ and ε_260_ values are implemented in the mRNACalc webserver. The source of these values is provided in the supplementary section. **b**, Steady-state absorption spectra of cytidine (black line) and 5-methylcytidine (red line) at pH 7.4. **b**, Steady-state absorption spectra of pseudouridine (orange line) and N1-methylpseudouridine (green line) at pH 7.4. **d**, Steady-state absorption spectra of uridine (light blue line) and N1-methylpseudouridine (green line) at pH 7.4. The ε_260_ for U and m^1^Ψ are shown. **e**, Relative UV absorption curves from mRNAs with uridine or N1-methylpseudouridine nucleosides. They were normalized to the corresponding F507 values and plotted relative to the peak maximum (ε_max_) of the U mRNA. **f**, The relative A_260_/F_507_ values from five replicates of the U-, Ψ-, and m^1^Ψ-mRNAs are shown, the black line corresponds to the average absorbance. Values are relative to the average absorbance of the U-mRNA. The three comparisons of the mean relative A_260_/F_507_ values were significant (t-test; p< 0.005).

To assess whether the complete U-to-m^1^Ψ substitution alter the UV absorbance of an mRNA, three mRNAs were transcribed using either U, Ψ, or m^1^Ψ. These mRNA also encoded a double-Broccoli aptamer in their 3’ untranslated region. Once the DFHBI-1T fluorophore was bound to the G-quadruplex in the Broccoli aptamer, the mRNA emitted green light upon excitation (15). The brightness, melting point and affinity of the DFHBI-1T-Broccoli complex were not significantly perturbed by the U-to-Ψ or U-to-m^1^Ψ substitutions (Supplementary Table 1 and Supplementary Figure 1). By normalizing the UV absorbance (A_260_) of each mRNA by its corresponding fluorescence (F_507_), it was observed that in practice the relative UV absorbance of the nucleoside-modified mRNA was significantly reduced as compared to the standard mRNA (ΔA_260_ = -10.6 %, Fig. 1e and 1f). The hypochromicity was slightly more pronounced in a second m^1^Ψ-mRNA with higher m^1^Ψ composition (ΔA_260_ = -11.8%, Supplementary figure 2). In principle, the modified nucleosides may also promote mRNA folding and reduce its UV absorption. This is particularly relevant for the pseudouridine modification. Its N1-hydrogen can engage in additional hydrogen bonds, promoting and stabilizing RNA folding. For instance, the U-to-Ψ substitution in tRNA stabilize the folded structure that is essential for translation (reviewed in ref. 16). However, the m^1^Ψ nucleobase lacks this additional hydrogen bonding capability, and it is expected to have little or no effect on the RNA folding of low CG-content (< 45 %) and less-structured RNA molecules, such as our mRNAs. Considering that both Ψ- and m^1^Ψ-mRNAs followed the anticipated hypochromicity associated to the nucleoside hypochromicity at 260 nm wavelength, instead of their expected distinct contribution to RNA folding, these data suggest that the observed reduction in nucleoside-modified mRNA UV absorption is mainly determined by the nucleobase composition and the intrinsic MAC of the nucleosides in these mRNA. Importantly, the UV absorption spectrum of the m^1^Ψ mRNA also depicted a broad absorption peak and a red shift, which brings additional implications for the assessment of the RNA sample purity (Supplementary note). These findings indicate that, for accurate determination of nucleoside-modified mRNA concentrations and proper interpretation of dose-ranging preclinical studies, the reported UV spectroscopic changes must be accounted for to prevent under-estimating mRNA concentrations by 5 to 15%, depending on the proportion of m^1^Ψ in the mRNA composition.

**Fig. 2:**
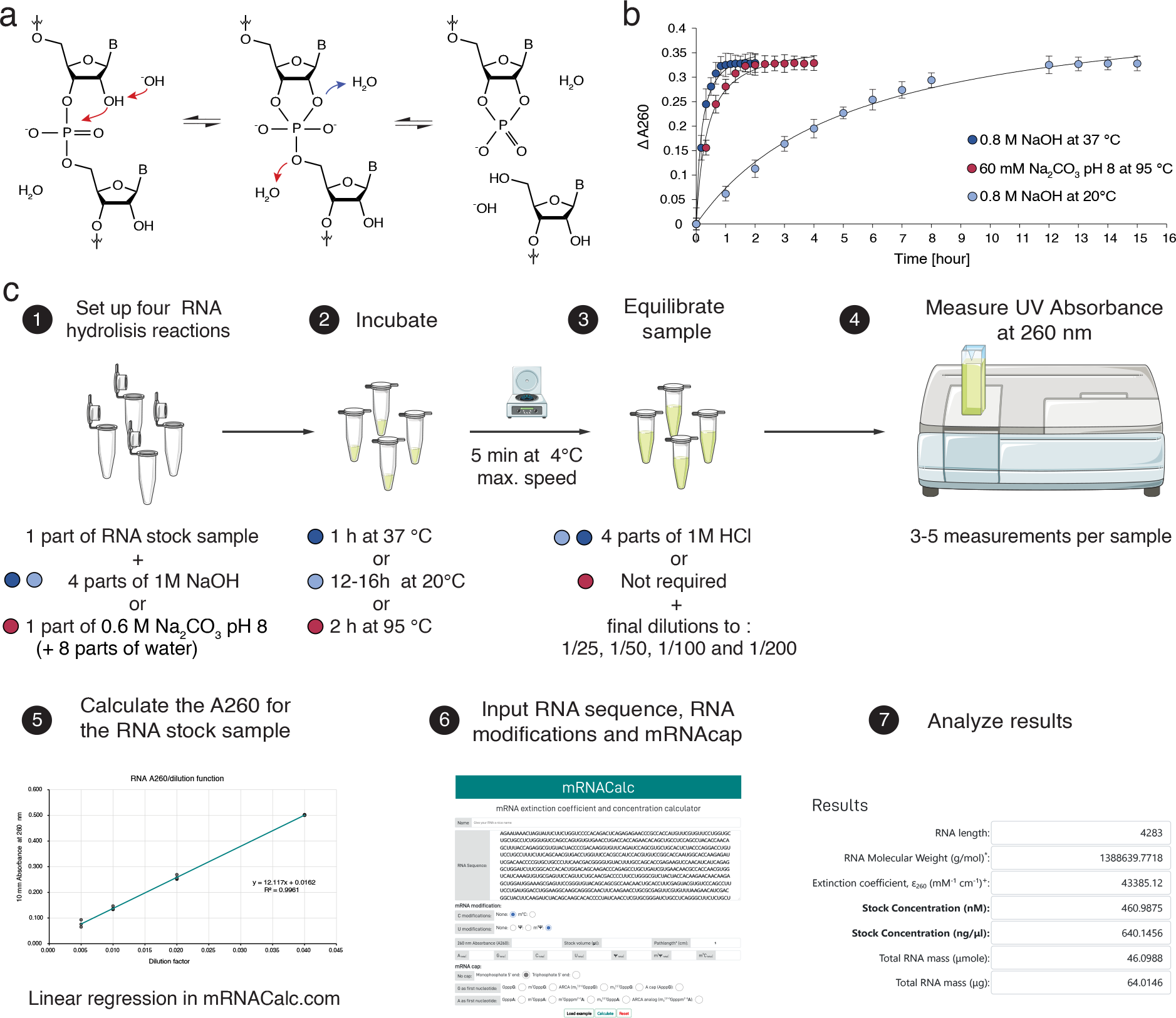
The mRNACalc webserver and the photometric measurement of RNA concentrations rely on the intrinsic UV molar absorption properties of nucleotides. **a**, Alkali-promoted transesterification allows RNA hydrolysis and mRNA quantification. Under alkaline conditions, the reactive -OH triggers the nucleophilic attack of the 2’-OH on the 3’,5’-phosphodiester linkage, converting the ground-state configuration of RNA into a penta-coordinated intermediate and leading to a 2’3’-cyclic phosphodiester. This cyclic form is then known to form 3’ and 2’ monophosphate nucleotides (not shown). **b**, Thermal and/or alkaline hydrolysis of RNA over time. Yeast RNA was hydrolysed using three different previously described methods and the ΔA_260_ was determined using an UV spectrophotometer at different intervals. For expedite RNA hydrolysis (1-or 2-hours incubation), a combination of thermal and alkaline hydrolysis can be used (dark blue dots, 0.8 M NaOH at 37 °C; red dots, 0.5 M Na_2_CO_3_ pH 8 at 95 °C). For overnight incubation, alkaline hydrolysis suffices (light blue dots, 0.8 M NaOH at 20 °C, the last four measurements were performed after an overnight incubation). Dots indicate the mean value of three measurements. Error bars correspond to standard deviations. **c**, Experimental workflow for the determination of RNA concentration using the mRNACalc web server. The coloured dots refer to the different RNA hydrolysis methods in 2.b.

To enable accurate measurement of nucleoside-modified mRNA, we also assessed different RNA hydrolysis and built an experimental workflow for the determination of mRNA concentrations. The modern analytical use of alkaline hydrolysis of RNA is known since 1922, when Steudel and Peiser demonstrated that 1 M NaOH hydrolysed yeast RNA, whereas thymus DNA resisted the NaOH hydrolysis (17). The alkali-promoted transesterification of RNA occurs due to the nucleophilic attack of the 2’-OH in the ribose to the 3’,5’-phosphodiester bond, explaining the alkali-resistance of the 2’-deoxyribonucleotides (Fig. 2a) (18). This reaction is further catalysed with the introduction of heat. However, the combination of thermal and alkaline hydrolysis, e.g., 1 M NaOH at 95 °C, also catalyses the deamination of cytosine to uridine in a small percentage of residues (19, 20). Thus, a compromise between the two methods is often applied. In our hands, three of such protocols showed a similar change in A_260_ upon hydrolysis of yeast RNA – a historical standard sample for these methods. (Fig. 2b). These protocols are provided in the mRNACalc web server along with a workflow that implement a linear regression model from multiple measurements at serial dilutions to reduce the impact of sample handling variation (Fig. 2c).

In the mRNACalc web server, the MACs of distinct modified nucleosides that form the capping nucleotide in the 5’ mRNA cap were also implemented for the sake of completeness. The capping nucleotide only represents one nucleotide out of thousands of nucleotides in a mRNA molecule and its contribution to the mRNA molar absorption is rather negligible.

Taken together, by incorporating modified nucleoside MAC parameters, accounting for the actual mRNA nucleotide composition, and providing an experimental workflow, the mRNACalc web server represents a freely available and all-inclusive tool for the determination of nucleoside-modified mRNA concentrations using UV spectroscopy.

## Materials and Methods

### The Beer-Lambert experiments

Pseudouridine (≥ 98% purity), 5-methylcytidine (≥ 99% purity), Cytidine (99% purity) and Uridine (99% purity) were purchased from Sigma-Aldrich. N1-methylpseudouridine (>95% purity) was purchased from Biosynth Carbosynth. They were used as received. Phosphate buffer solutions with a total phosphate concentration of 16 mM from monosodium and disodium phosphate salts dissociated in ultrapure water (Milli-pore) were freshly prepared on the day of each experiment. The pH of the solution was adjusted using 0.1 M solutions of NaOH and HCl to the desired pH of 7.4 (± 0.1 pH units). Steady-state absorption was recorded using a Cary 100 spectrometer. Serial dilutions of known concentration were carried out such that the absorbance reading at the respective lambda maximum (local maximum absorbance) remained below 1.0, within the linear range of the instrument. The molar absorption coefficients were experimentally determined using the slope from the linear regression from plotting absorbance versus concentration. The correlation constant for the linear regression analysis of the Beer-Lambert’s Law data for determining molar absorption constants was >0.9999 showing a strong linear relationship.

### mRNA *in vitro* transcription and purification

The Plasmid DNA template (pUCIDT plasmid) was grown in DH5 alpha *E. coli* (New England Biolabs, Inc.) in 300 ml of Luria-Bertani broth supplemented with Kanamycin (50 μg/ml) and a maxi preparation was performed using the QIAGEN® Plasmid Plus Maxi Kit following manufacturer instructions. The plasmid encoded a T7 promoter followed by the mCherry gene with a degradation tag (1449 nucleotides) plus the 3’ and 5’ untranslated regions (UTR) of the BNT162b2 mRNA vaccine (541 nucleotides). The double broccoli aptamer was encoded within the poli-adenine region in the 3’UTR. The plasmid was linearized by EcoRV restriction enzyme digestion at the end of the 3’ UTR.

A standard T7 transcription reaction included 30 mM Tris-HCl, pH 7.9, 2 mM spermidine, 30 mM MgCl_2_, 5 mM NaCl, 10 mM DTT, 50 μg/ml BSA (New England Biolabs, Inc.), 0.005% Triton X-100, 2% polyethylene glycol (PEG8000), 5 mM of each triphosphate ribonucleotide (standard nucleotides were purchased from Jena Bioscience GmBH and pseudouridine and N1-methylpseudouridine from BOC sciences), 2 μM linearized plasmid DNA template, 3.5 μM T7 RNA Polymerase (in house produced and purified) and 0.0025 units of *E. coli* inorganic PPase (New England Biolabs, Inc). All reagents were purchased from Sigma-Aldrich, unless otherwise stated. The reactions were incubated at 37 °C for 2.5 hours and stopped by the addition of 500 mM EDTA, pH 8 to a final concentration of 35 mM.

The mRNA was purified using anion exchange chromatography. A PRP-X600 Anion exchange column (Hamilton Company, Inc.) was equilibrated in Buffer A (85:15 100 mM TRIS, pH 8/Acetonitrile). RNA samples were loaded onto the column at a flow rate of 3 ml/min and eluted with a 40-minutes gradient of 0-40% buffer B (85:15 100 mM TRIS 2.5 M LiCl, pH 8/Acetonitrile). Fractions containing the mRNA were collected and the mRNA molecules were precipitated using standard Butanol extraction (21). The purity of the mRNA preparation was assessed using high-resolution automated electrophoresis in the Agilent 2100 Bioanalyzer system using the Bioanalyzer RNA 6000 pico assay (Agilent Technologies, Inc).

### Determination of the mRNA UV absorption spectrum

To determine the UV absorption spectrum of mRNAs, the mRNAs stocks were diluted to approximately 25 nM into a buffer containing 40 mM HEPES pH 7.4, 5 mM MgCl_2,_ and 100 mM KCl to a final volume of 2 ml. Five independent mRNA samples were prepared per mRNA set (U-, Ψ-, and m^1^Ψ-mRNA). The UV absorption spectra were recorded for each mRNA sample using in a UV-3600i plus UV-VIS spectrophotometer (Shimadzu Corp.).

### Excitation-emission experiments on the DFHBI-1T bound mRNAs

After UV absorption determination, the mRNA samples were bound to the DFHBI-1T fluorophore, by adding 100 μM DFHBI-1T, 100% DMSO to a 500 nM concentration into the 2-ml mRNA samples. Fluorescence was measured using a Fluorolog-3 spectrofluorometer (Horiba Scientific) using the excitation and emission wavelengths commonly used for DFHBI-1T (Excitation: 472 nm, emission: 507 nm) (15).

### Determination of the relative UV absorbance (A_260_)

The A_260_/F_507_ ratios were calculated for each mRNA sample. The mean A_260_/F_507_ values for U-, Ψ-, and m^1^Ψ-mRNA were calculated. The A_260_/F_507_ values of each sample were normalized using the mean A_260_/F_507_ value from the U-mRNA as reference and they were plotted in a dot plot. The t-tests were applied to compare the mean A_260_/F_507_ values across each pair of mRNA sets, using a p-values of 0.005 as cut-off of significance.

### Methods of RNA hydrolysis

Two methods of RNA hydrolysis were tested in this study. Torula yeast RNA was used as standard RNA sample (Sigma-Aldrich). The Yeast RNA stock was prepared at 1000 μg/μl in water. Thus, after 1/25 dilution, the UV absorbance of this RNA sample would be within the linear range of the instrument (UV-3600i plus UV-VIS spectrophotometer, Shimadzu Corp.).

The most extensively used alkaline RNA hydrolysis method involves adding 1 part of RNA and 4 parts of 1 M NaOH and incubating them at 37 °C for 1 hour (22). To test this method, twelve yeast RNA samples were hydrolysed. Every 10 minutes, a sample was neutralised with 4 parts of 1 M HCl and diluted to 1/25 with 16 parts of water. Three UV absorbance measurements were performed on every sample. Similarly, a room temperature variation of this method is often used for overnight RNA hydrolysis. Therefore, twelve RNA samples were hydrolysed and incubated at 20 °C for up to 15 hours. Samples were neutralized and diluted hourly followed by three UV absorbance measurements.

A second method of thermal hydrolysis at neutral pH was also tested (8). To test this method, twelve yeast RNA samples hydrolysed (1 part of RNA in 9 parts of 60 mM Na_2_CO_3_ pH 8) with an incubation of at 95 °C for up to 2 hours. Every 20 minutes, a sample was diluted to 1/25 with 15 parts of water and three UV absorption measurements were performed on every sample.

## Supporting information

Supplementary section

## Code availability

The webserver is available at https://www.mrnacalc.com. The website is free and open to all users and there is no login requirement.

The source code for mRNACalc webserver is available under GNU general public licence from https://github.com/estebanfbfc/mRNACalc.

## Contributions

E.F. conceived the study. E.F. and C.E.C-H. supervised the project. E.F. developed the mRNACalc webserver. S.E.K. and S.J.H. performed the Beer-Lambert experiments and prepared the corresponding figure panel. E.F. performed the relative absorbance of mRNA experiments and analysed the data. E.F. prepared figures, wrote the initial draft of the manuscript and edited the submitted version of the manuscript with contributions from all the authors.

## Acknowledgement

The authors would also like to thank Prof. Eng Eong Ooi for his invaluable advice and generosity throughout this work and Prof. Guillermo C. Bazan for providing access to the UV-Vis and fluorescence spectrometers in his laboratory.

## Funding

This work was supported by the National Medical Research Council through an Open Fund - Large Collaborative Grant, granted to Prof. Eng Eong Ooi, and by the National Science Foundation (Grant No. CHE-2246805), granted to Prof. Carlos E. Crespo-Hernández.

## Conflict of interest statement

Authors declare no competing interests.

## Notes

### Competing Interest Statement

The authors have declared no competing interest.

### Summary of Updates

Results from additional experiments were added.

## References

1. Chaudhary, N., Weissman, D. and Whitehead, K.A. (2021) mRNA vaccines for infectious diseases: principles, delivery and clinical translation. Nat Rev Drug Discov, 20, 817–838.

2. Sahin, U., Karikó, K. and Türeci,Ö. (2014) mRNA-based therapeutics — developing a new class of drugs. Nat Rev Drug Discov, 13, 759–780.

3. Webb, C., Ip, S., Bathula, N.V., Popova, P., Soriano, S.K.V., Ly, H.H., Eryilmaz, B., Nguyen Huu, V.A., Broadhead, R., Rabel, M., et al. (2022) Current Status and Future Perspectives on MRNA Drug Manufacturing. Mol Pharm, 10.1021/acs.molpharmaceut.2c00010.

4. Karikó, K., Muramatsu, H., Welsh, F.A., Ludwig, J., Kato, H., Akira, S. and Weissman, D. (2008) Incorporation of Pseudouridine Into mRNA Yields Superior Nonimmunogenic Vector With Increased Translational Capacity and Biological Stability. Molecular Therapy, 16, 1833–1840.

5. Andries, O., Mc Cafferty, S., De Smedt, S.C., Weiss, R., Sanders, N.N. and Kitada, T. (2015) N1-methylpseudouridine-incorporated mRNA outperforms pseudouridine-incorporated mRNA by providing enhanced protein expression and reduced immunogenicity in mammalian cell lines and mice. Journal of Controlled Release, 217, 337–344.

6. Cavaluzzi, M.J. and Borer, P.N. (2004) Revised UV extinction coefficients for nucleoside-5′-monophosphates and unpaired DNA and RNA. Nucleic Acids Res, 32, e13.

7. Tinoco, I.Jr. (1960) Hypochromism in Polynucleotides1. J. Am. Chem. Soc., 82, 4785–4790.

8. Wilson, S.C., Cohen, D.T., Wang, X.C. and Hammond, M.C. (2014) A neutral pH thermal hydrolysis method for quantification of structured RNAs. RNA, 20, 1153–1160.

9. Doty, P., Boedtker, H., Fresco, J.R., Haselkorn, R. and Litt, M. (1959) Secondary structure in ribonucleic acids*. Proceedings of the National Academy of Sciences, 45, 482–499.

10. Cohn, W.E. (1960) Pseudouridine, a Carbon-Carbon Linked Ribonucleoside in Ribonucleic Acids: Isolation, Structure, and Chemical Characteristics. Journal of Biological Chemistry, 235, 1488–1498.

11. Stimson, M.Michael. (1949) The Ultraviolet Absorption Spectra of Some Pyrimidines. Chemical Structure and the Effect of pH on the Position of λmax. J. Am. Chem. Soc., 71, 1470–1474.

12. Sharonov, A., Gustavsson, T., Marguet, S. and Markovitsi, D. (2003) Photophysical properties of 5-methylcytidine. Photochem Photobiol Sci, 2, 362–364.

13. Shugar, D. and Fox, J.J. (1952) Spectrophotometric studies op nucleic acid derivatives and related compounds as a function of pH: I. Pyrimidines. Biochimica et Biophysica Acta, 9, 199–218.

14. Rabczenko, A. and Shugar, D. (1971) Studies on the conformation of nucleosides, dinucleoside monophosphates and homopolynucleotides containing uracil or thymine base residues, and ribose, deoxyribose or 2’-O-methylribose. Acta Biochim Pol, 18, 387–402.

15. Filonov, G.S., Moon, J.D., Svensen, N. and Jaffrey, S.R. (2014) Broccoli: Rapid Selection of an RNA Mimic of Green Fluorescent Protein by Fluorescence-Based Selection and Directed Evolution. J. Am. Chem. Soc., 136, 16299–16308.

16. Lorenz, C., Lünse, C.E. and Mörl, M. (2017) tRNA Modifications: Impact on Structure and Thermal Adaptation. Biomolecules, 7, 35.

17. Steudel, H. and Peiser, E. (1922) Über Nucleinsäure-Eiweißverbindungen. 122, 298–306.

18. Lipkin, D., Talbert, P.T. and Cohn, M. (1954) The Mechanism of the Alkaline Hydrolysis of Ribonucleic Acids. J. Am. Chem. Soc., 76, 2871–2872.

19. Wang, R.Y.-H., Kuo, K.C., Gehrke, C.W., Huang, L.-H. and Ehrlich, M. (1982) Heat- and alkali-induced deamination of 5-methylcytosine and cytosine residues in DNA. Biochimica et Biophysica Acta (BBA) - Gene Structure and Expression, 697, 371–377.

20. Shen, J.C., Rideout, W.M. and Jones, P.A. (1994) The rate of hydrolytic deamination of 5-methylcytosine in double-stranded DNA. Nucleic Acids Res, 22, 972–976.

21. Green, M.R. and Sambrook, J. (2017) Concentrating Nucleic Acids by Extraction with Butanol. Cold Spring Harb Protoc, 2017, pdb.prot093401.

22. Bock, R.M. (1967) [29] Alkaline hydrolysis of RNA. In Methods in Enzymology, Nucleic Acids, Part A. Academic Press, Vol. 12, pp. 224–228.

