## Supplementary section for "The mRNACalc web server accounts for the hypochromicity of modified nucleosides and enables the accurate quantification of nucleoside-modified mRNA"

### Supplementary data:

**Supplementary Table 1: photophysical and biochemical properties of mutated Broccoli−DFHBI-1T complexes.**

| Complex | $\lambda_{max}$ (nm) | Relative brightness* | $K_D$ (nM)$^+$ | $T_m$ (°C)$^+$ |
|---|---|---|---|---|
| U-Broc−DFHBI-1T (ref. 15) | 507 | ------ | 360 | 48 |
| U-Broc−DFHBI-1T | 507 | 1.000 ± 0.002 | 379.6 ± 13.89 | 49.13 ± 0.13 |
| Ψ-Broc-DFHBI-1T | 507 | 1.005 ± 0.004 | 378.7 ± 8.11 | 49.46 ± 0.09 |
| m$^1$Ψ-Broc-DFHBI-1T | 507 | 1.004 ± 0.003 | 375.6 ± 8.17 | 49.23 ± 0.07 |

*Relative to the U-Broc-DFHBI-1T complex. Data are shown as mean ± SD.

$^+$ Data are shown as fitted $K_D$ ± Error of the fit or fitted $T_m$ ± Error of the fit.

**Supplementary Figure 1: binding and melting curves of mutated Broccoli−DFHBI-1T complexes.**

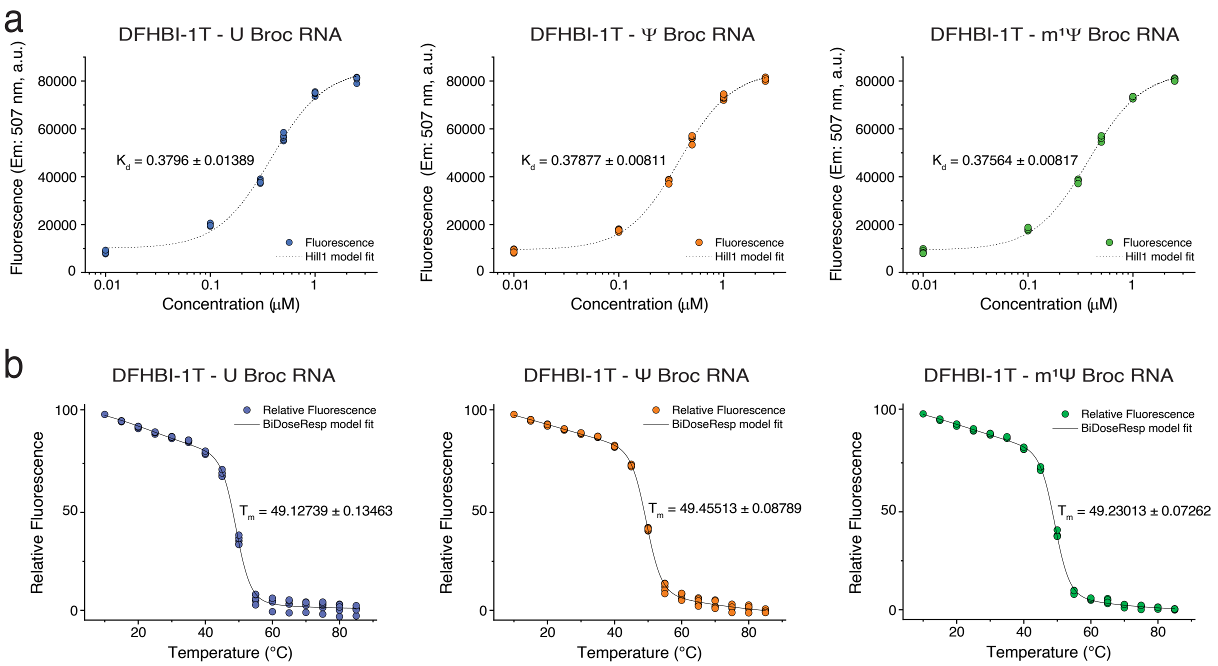


**a**, the binding curves of DFHBI-1T onto the U-, Ψ- and m$^1$Ψ-broccoli RNA aptamers are shown.
**b**, the melting curves of DFHBI-1T onto the U-, Ψ- and m$^1$Ψ-broccoli RNA aptamers are shown.

**Supplementary Figure 2: The nucleotide composition of mRNA determines their UV absorption at 260 nm wavelength**.

The relative A260/F507 values from five replicates of two different mRNAs bearing either U or $m^1\Psi$ nucleosides are shown. The data from mRNA1 is also shown in Figure 1e in the main text. The nucleotide composition of the two mRNAs is shown at the bottom of the graph. The black line corresponds to the average absorbance. Values are relative to the average absorbance of the U-mRNAs. The comparisons of the mean relative A260/F507 values were significant (t-test; $p< 0.005$).

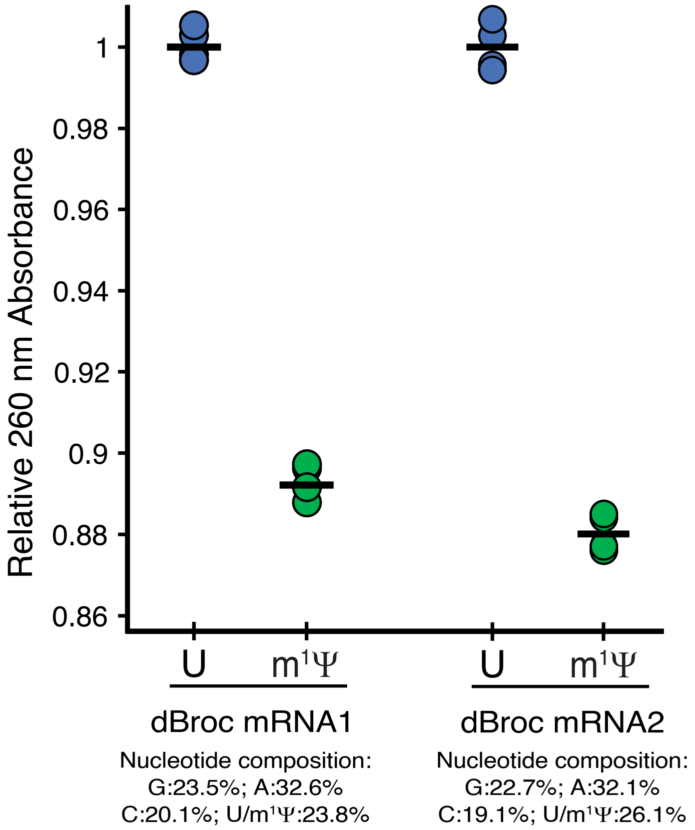


### Methods for supplementary data:

#### Determination of photophysical and biochemical properties of mutated Broccoli−DFHBI-1T Complexes

The $\lambda_{max}$, relative brightness, dissociation constants and melting points were determined following the methods in reference 15 of the main text.

The emission was measured for solutions using “excess RNA” conditions, to ensure that no free fluorophore contributes to the fluorescence signal. The RNA concentration was 30 µM, while DFHBI-1T concentration was 2 µM. The fluorescence emission was determined in a Fluorolog-3 spectrofluorometer (Horiba Scientific) using the excitation wavelength commonly used for DFHBI-1T, 472 nm, with a side entrance and side exit slits of 3 nm. The integration time was 0.1 seconds, the emission was recorded from 482 nm to 700 nm, with 1 nm increments. The side entrance, front exit and side exit slits were 3 nm.

For the relative brightness, the fluorescence signal of Broccoli-DFHBI-1T complex was compared at different dilutions, using the U-Broccoli-DFHBI-1T as reference.

To calculate dissociation constant ($K_D$), we titrated increasing concentrations of DFHBI-1T into 50 nM of RNA. The fluorescence at 507 nm wavelength was determined in a Fluorolog-3 spectrofluorometer (Horiba Scientific), using the excitation wavelengths commonly used for DFHBI-1T (Excitation: 472 nm). The resulting data points were fitted to the Hill equation using Origin Pro Software.

To measure the thermostability of RNA-fluorophore complexes, 50 nM of RNA were incubated with 300 µM DFHBI-1T. Then fluorescence values were recorded in 5 °C increments from 10

°C to 85 °C, with a 2-min incubation at each temperature to allow for equilibration, using a CFX96 thermocycler (Bio-rad). The resulting data points were fitted to a biphasic model using Origin Pro Software.

**Determination of A260/F507 ratio on dBroc mRNAs:**

These methods are described in the Material and Methods section.

**Supplementary notes:**

**On the purity of RNA samples:**

The presence of impurities in nucleic acid samples is often assessed using the $A_{260:280}$ and $A_{260:230}$ ratios. For pure RNA the $A_{260:280}$ ratio is ~ 2.0. This ratio is commonly used to assess the amount of protein contamination, since proteins absorb at 280 nm. Similarly, the $A_{260:230}$ ratio for pure RNA is often slightly higher than the $A_{260:280}$ ratio, ranging from 2.0 to 2.2. Residual chemical contamination (phenol, butanol, carbohydrates, guanidine, and others) from the RNA purification method can increase the $A_{230}$ and reduce the $A_{260:230}$ ratio.

From our experience, the assessment of purity for the $m^1\Psi$ modified mRNA samples requires shifting the wavelength for these ratios to $A_{264:284}$ and $A_{264:234}$ due to the bathochromic shift in the mRNA absorption curve. Which leads to:

- a reduced $A_{260}$ due to the $\lambda_{max}$ shift ($\lambda_{max}$ at ~264 nm),
- an increased $A_{230}$ due to the shift on the curve trough ($\lambda_{min}$) to ~234 nm,
- and an increased $A_{280}$ due to the absorbance peak shift and broadening. The broadening arises due to the wider range of $\lambda_{max}$ values in the RNA composition (standard mRNA= 252 to 263 nm, $m^1\Psi$ modified mRNA= 252 to 272 nm) as well as, due to the broader absorption peak of $m^1\Psi$ (as determined by the peak width at the trough level: Urd= 52.5 nm vs $m^1\Psi$= 57.0 nm).

Thus, $A_{264:284}$ and $A_{264:234}$ ratios should be interpreted in the same manner as the $A_{260:280}$ and $A_{260:230}$ ratios, respectively. Alternatively, the $A_{260:280}$ ratio can be accepted at 1.9 and the $A_{260:220}$ ratio can range between 1.9 to 2.1.

Importantly, $m^5C$-modified mRNA should show similar modifications in the UV molar absorption spectrum and the proposed shifted ratios may be applied as well.

**On the mRNAcalc web server calculations**

- The mRNA molar absorption coefficient ($\varepsilon$) is calculated from the sum of the individual nucleotide extinction coefficients as determined by:

$$\varepsilon_{mRNA} = n_A\varepsilon_A + n_G\varepsilon_G + n_C\varepsilon_C \; or \; n_{m^5C}\varepsilon_{m^5C} + n_U\varepsilon_U \; or \; n_\Psi\varepsilon_\Psi \; or \; n_{m^1\Psi}\varepsilon_{m^1\Psi} + \varepsilon_{cap}$$

Where $n_N$ corresponds to the number of each type of nucleotide, N, in the mRNA and $\varepsilon_N$ to the molar absorption coefficient for each type of nucleotide, including the capping nucleotide ($\varepsilon_{cap}$).

- The mRNA molecular weight is calculated as the sum of the nucleotide composition mass as RNA-incorporated monophosphate nucleotides.

$$MW_{mRNA} = n_A MW_A + n_G MW_G + n_C MW_C \; or \; n_{m^5C} MW_{m^5C} + n_U MW_U \; or \; n_\Psi MW_\Psi \; or \; n_{m^1\Psi} MW_{m^1\Psi} + MW_{cap}$$

- The mRNA molar concentration is calculated using the Beer-Lambert equation:

$$Concentration\ (M) = \frac{A260}{\varepsilon_{mRNA} * cm^{-1}}$$

The mRNA molar concentration is presented in the nM and ng/µl scales in the webserver.

**On the molar absorption coefficients of nucleosides/nucleotides:**

For standard nucleotides, the mRNACalc web server implements the $\varepsilon_{260}$ parameters in the supplementary Table 2. The parameters from Cavaluzzi et al. were obtained after accurate measurements of nucleotides concentration using nuclear magnetic resonance spectroscopy.

**Supplementary Table 2: Molar absorption coefficients of standard nucleosides as reported in Cavaluzzi et al.**

| Standard nucleosides | $\lambda_{max}$ (nm) | $\varepsilon_{max}$ ($mM^{-1}cm^{-1}$) | $\varepsilon_{260}$ ($mM^{-1}cm^{-1}$) |
|---|---|---|---|
| Uridine | 262 | 9.78 | 9.66 |
| Uridine* | 262 | 9.66 | 9.60 |
| Thymidine | 267 | 9.49 | 8.56 |
| Cytidine | 271 | 8.74 | 7.07 |
| Cytidine* | 271 | 9.34 | 7.67 |
| Guanosine | 252 | 14.09 | 12.08 |
| Adenosine | 259 | 15.04 | 15.02 |

Source: Cavaluzzi et al. (1) and *this study.

For the modified nucleosides/nucleotides, we have determined and searched for $\varepsilon_{max}$ and $\varepsilon_{260}$ parameters in the literature and in the datasheet of $\Psi$, $m^1\Psi$, and $m^5C$ manufacturers, which are summarized in the Supplementary Tables 3, 4 and 5.

Considering the extensive variability across the published and manufacturer-provided values for $m^5C$ and $\Psi$, the mRNACalc webserver implements the average $\varepsilon_{260}$ values. For $m^1\Psi$, the mRNACalc server implements the $\varepsilon_{260}$ value that was obtained for this study, due to the limited number of previously reported values.

**Supplementary Table 3: Molar absorption coefficients of pseudouridine as reported in the literature and manufacturers' datasheets.**

| **Pseudouridine** | | | |
|---|---|---|---|
| Source | $\lambda_{max}$ (nm) | $\varepsilon_{max}$ ($mM^{-1}cm^{-1}$) | $\varepsilon_{260}$ ($mM^{-1}cm^{-1}$) |
| Basanta-Sanchez et al. (2) | 262 | 7.583 | 7.492 |
| Yu & Allen.(3) | 263 | 7.5 | Not provided |
| David & Allen. (4) | 263 | 8.4 | 8.3 |
| Shapiro & Chambers. (5) | 262 | 7.9 | Not provided |
| Michelson & Cohn. (6) | 262 | 8.0 | Not provided |
| Cohn. (7) | 263 | 8.1 | Not provided |
| Jena Biosciences | 265 | 7.9 | Not provided |
| Trilink Biotechnologies | 262 | 7.546 | Not provided |
| This study | 263 | 7.677 | 7.527 |
| Average | 263 | 7.877 | 7.723$^{+}$ |

$^{+}$Average $\varepsilon_{260}$ was calculated by multiplying the average $\varepsilon_{max}$ by the observed $\varepsilon_{260/263}$ ratio.

**Supplementary Table 4: Molar absorption coefficients of N1-methylpseudouridine as reported in the literature and manufacturers' datasheets.**

| **N1-methylpseudouridine** | | | |
|---|---|---|---|
| Source | $\lambda_{max}$ (nm) | $\varepsilon_{max}$ ($mM^{-1}cm^{-1}$) | $\varepsilon_{260}$ ($mM^{-1}cm^{-1}$) |
| Roche | 271 | 7.3 | Not provided |
| Trilink Biotechnologies | 271 | 8.877 | Not provided |
| This study | 272 | 7.726 | 5.813 |
| Average | 271 | 7.967 | 5.994$^{+}$ |

$^{+}$Average $\varepsilon_{260}$ was calculated by multiplying the average $\varepsilon_{max}$ by the observed $\varepsilon_{260/272}$ ratio.

**Supplementary Table 5: Molar absorption coefficients of N1-methylpseudouridine as reported in the literature and manufacturers' datasheets.**

| **5-methylcytidine** | | | |
|---|---|---|---|
| Source | $\lambda_{max}$ (nm) | $\varepsilon_{max}$ ($mM^{-1}cm^{-1}$) | $\varepsilon_{260}$ ($mM^{-1}cm^{-1}$) |
| Szer. (8) | 278.5 | 8.8 | Not provided |
| Martínez-Fernández et al.(9) | 278 | 8.92 | Not provided |
| Fox et al.(10) | 277.5 | 8.88 | Not provided |
| Ma et al.(11) | 278 | 8.871 | Not provided |
| Fox & Shugar. (12) | 276 | 8.05 | Not provided |
| Shanorov et al.(13) | 278 | 8.4 | Not provided |
| Jena Biosciences | 277 | 9 | Not provided |
| Sigma-Aldrich | 278 | 8.5 | Not provided |
| Glenn research | 277 | 9 | Not provided |
| Trilink Biotechnologies | 279 | 7.808 | Not provided |
| This study | 278 | 7.948 | 5.199 |
| Average | 278 | 8.562 | 5.601$^{+}$ |

$^{+}$Average $\varepsilon_{260}$ was calculated by multiplying the average $\varepsilon_{max}$ by the observed $\varepsilon_{260/278}$ ratio.

For the mRNA capping nucleotides, the mRNACalc web server implements the $\varepsilon_{260}$ values provided by the manufactures (Supplementary Table 6). In few cases, the $\varepsilon_{260}$ values were not available, the independent $\varepsilon_{260}$ values of the two nucleotides were summed up. Overall, these mRNA cap parameters were only implemented for completeness, and they can be considered as rough estimations, despite their contribution to an mRNA UV absorption is rather negligible.

**Supplementary Table 6: Molar absorption coefficients of mRNA capping nucleotides as reported in the manufacturers' datasheets.**

| mRNA cap | $\varepsilon_{260}$ ($mM^{-1}cm^{-1}$) |
|---|---|
| GpppG | 24.16 |
| $m^7$GpppG | 22.31 |
| ARCA ($m_2^{7,3'-O}$GpppG) | 20.46 |
| $m_3^{2.2.7}$GpppG | 21.6 |
| ApppG | 27.1 |
| GpppA | 27.1 |
| $m^7$GpppA | 25.25 |
| m7Gppp $m^{2'-O}$A | 23.43 |
| $m_3^{2.2.7}$GpppA | 24.54 |
| ARCA analog ($m_2^{7,3'-O}$Gpppm$^{2'-O}$A) | 20.28 |

Important note: The molar absorption parameters, herein compiled, correspond to either nucleosides or nucleotides in aqueous buffered solution (pH 7 – 8). Considering that the contribution of the phosphate group to the molar absorption of nucleotides is negligible, they have been considered for their implementation in the mRNACalc web server indiscriminately.

References:


1. Cavaluzzi, M.J. and Borer, P.N. (2004) Revised UV extinction coefficients for nucleoside-5′-monophosphates and unpaired DNA and RNA. *Nucleic Acids Res*, **32**, e13.

2. Basanta-Sanchez, M., Temple, S., Ansari, S.A., D'Amico, A. and Agris, P.F. (2016) Attomole quantification and global profile of RNA modifications: Epitranscriptome of human neural stem cells. *Nucleic Acids Research*, **44**, e26.

3. Yu, C.-T. and Allen, F.W. (1959) Studies of an isomer of uridine isolated from ribonucleic acids. *Biochimica et Biophysica Acta*, **32**, 393–406.

4. Davis, F.F. and Allen, F.W. (1957) Ribonucleic acids from yeast which contain a fifth nucleotide. *Journal of Biological Chemistry*, **227**, 907–915.

5. Shapiro, R. and Chambers, R.W. (1961) synthesis of pseudouridine. *J. Am. Chem. Soc.*, **83**, 3920–3921.

6. Michelson, A.M. and Cohn, W.E. (1962) Cyclo-pseudouridine and the Configuration of Pseudouridine. *Biochemistry*, **1**, 490–495.

7. Cohn, W.E. (1960) Pseudouridine, a Carbon-Carbon Linked Ribonucleoside in Ribonucleic Acids: Isolation, Structure, and Chemical Characteristics. *Journal of Biological Chemistry*, **235**, 1488–1498.

8. Szer, W. (1965) Secondary structure of poly-5-methylcytidylic acid. *Biochemical and Biophysical Research Communications*, **20**, 182–186.

9. Martínez-Fernández, L., Pepino, A.J., Segarra-Martí, J., Banyasz, A., Garavelli, M. and Improta, R. (2016) Computing the Absorption and Emission Spectra of 5-Methylcytidine in Different Solvents: A Test-Case for Different Solvation Models. *J. Chem. Theory Comput.*, **12**, 4430–4439.

10. Fox, J.J., Van Praag, D., Wempen, I., Doerr ,I.L., Cheong, L., Knoll, J.E., Eidinoff, M.L., Bendich,A. and Brown,G.B. (1959) Thiation of Nucleosides. II. Synthesis of 5-Methyl-2'-deoxycytidine and Related Pyrimidine Nucleosides. *J. Am. Chem. Soc.*, **81**, 178–187.

11. Ma, C., Cheng, C.C.-W., Chan, C.T.-L., Chan, R.C.-T. and Kwok, W.-M. (2015) Remarkable effects of solvent and substitution on the photo-dynamics of cytosine: a femtosecond broadband time-resolved fluorescence and transient absorption study. *Phys. Chem. Chem. Phys.*, **17**, 19045–19057.

12. Shugar, D. and Fox, J.J. (1952) Spectrophotometric studies of nucleic acid derivatives and related compounds as a function of pH: I. Pyrimidines. *Biochimica et Biophysica Acta*, **9**, 199–218.

13. Sharonov, A., Gustavsson, T., Marguet, S. and Markovitsi, D. (2003) Photophysical properties of 5-methylcytidine. *Photochem Photobiol Sci*, **2**, 362–364.
